# Phylourny: Predicting the Knock-out-phase of Tournaments via Phylogenetic Methods by example of the UEFA EURO 2020

**DOI:** 10.1101/2021.06.24.449715

**Authors:** Ben Bettisworth, Alexandros Stamatakis

## Abstract

The prediction of knock-out tournaments represents an area of large public interest and active academic as well as industrial research. Here, we leverage the computational analogies between calculating the so-called phylogenetic likelihood score used in the area of molecular evolution and efficiently calculating, instead of approximating via simulations, the *exact* per-team winning probabilities, given a pairwise win probability matrix *P*. We implement and make available our method as open-source code and deploy it to calculate the winning probabilities for all teams participating at the knock-out phase of the UEFA EURO 2020 football tournament. We use three different *P* matrices to conduct predictions, two inferred via our own simple method and one computed by experts in the field. According to this expert *P* matrix which we trust most, we find that the most probable final is France versus England and that England has a slightly higher probability to win the title. The ability to efficiently and exactly compute winning probabilities, apart from improving and accelerating predictions, might allow for the development of novel methods to compute *P*.

## 1 Introduction

Predicting the winner of knock-out (bracket-based/elimination) tournaments can become computationally expensive if a high degree of accuracy shall be attained. To fully (and naïvely) evaluate the probability of the final placing of any particular tournament competitor, a polynomial with a comparatively large number of terms must be evaluated (see 3.2 for details). More specifically, for a tournament with *n* teams, a polynomial with 2*^n^* terms must be evaluated. If one desires to calculate this for every tournament competitor, then *n* such polynomials must be evaluated. Alternatively, one can use stochastic simulations in practice to estimate the probability distribution of the tournament winners. This can potentially be computationally more efficient, but comes at the cost of reduced fidelity of the results [1, 3].

However, there exists a similar problem in the field of computational phylogenetics, that is, the field of Bioinformatics that develops methods for reconstructing the evolutionary histories of currently living species based on their DNA or amino acid sequence data. Computational phylogenetics exhibits a plethora of similarities to the problem of computing the winning probabilities for a tournament. We will henceforth focus on the phylogenetic likelihood model [4] that is currently the most widely used model for phylogenetic inference (i.e., reconstructing evolutionary histories among extant species).

Initially, let us consider the problem of computing the likelihood score for a given statistical model of molecular sequence evolution on a given, possible evolutionary history (i.e., a phylogenetic tree). Note that, the specific phylogenetic tree whose likelihood shall be evaluated also constitutes a parameter of the likelihood model. This tree parameter is special in the sense that it represents the only *discrete* parameter of the phylogenetic likelihood model. While in general, phylogenetic trees are unrooted, without loss of generality for the purpose of knock-out tournament predictions, we can assume that they are rooted and hence *do* have a direction. In addition, for a tournament, the tree is already given which simplifies the task at hand. The likelihood score on a given tree topology under a given model can be efficiently computed using a dynamic programming algorithm called ‘Felsenstein pruning algorithm’ that was presented in Joe Felsenstein’s seminal paper that introduces the phylogenetic likelihood model [4].

The most striking similarity between the two problems is that computational phylogenetics and knock-out tournament predictions share a directed acyclic graph as a model parameter. Additionally, the shapes of these graphs are restricted in analogous ways, which allows to apply computational techniques from phylogenetics to tournament prediction.

In addition, both are based on statistical principles. Computational phylogenetics seeks to compute a likelihood, which is the probability of a model, given some data. So, while this is in principle different than ‘just’ computing a probability, the underlying structure and order of computations is highly similar. More importantly, the computation of the likelihood can be expressed via polynomials, following a procedure that is essentially analogous to computing the probability of a particular competitor winning a tournament. The above analogies allow us to adapt techniques which have been developed to efficiently compute phylogenetic likelihood scores to also efficiently compute tournament win probabilities.

In the following, we propose a novel method of computing win probabilities for a multi-elimination tournament^1^, which allows for the exact calculation of win probabilities in conjunction with high computational efficiency. Our novel method, which we call Phylourny, is based on an observation by Ziheng Yang [10] about the aforementioned Felsenstein pruning algorithm. Ziheng Yang points out that Felsenstein’s algorithm can be interpreted as an efficient way to compute polynomials of a high degree.

We implement our new method in a software tool that is also called Phylourny. The name is a portmanteau of Phylogeny and tournament. We show that methods which use a similar evaluation strategy as Phylourny are substantially faster than naïve tournament prediction approaches. We (will) also assess our method by predicting the winner and winning probabilities of the teams participating at the knock-out phase of the UEFA EURO 2020 European Football Championship^2^. As already mentioned, a pairwise win prediction matrix *P* is required as input for our method. We utilize a *P* matrix based on prior work by experts in the field and two *P* matrices obtained via a simple method implemented in Phylourny that solely uses match data from the group stage. Using these data sources, we obtain three predictions, which are summarized in Section 3.4.

## 2 Background

A graph is a set of nodes, and the relationships between those nodes, are called edges. A directed graph is a graph where the edges, here called arcs, have a direction. For example, in Figure 1 there is an arc from `a` to `n1`, but not vice versa. A directed acyclic graph (DAG) constitutes a special, simpler form of a directed graph that does not contain cycles. A graph *has* a cycle if starting from some node, there exists a set of edges which lead back to the same node. Alternatively, one can define a directed graph to be acyclic if, when some node *a* can be reached from *b* this implies that *b* can not be reached from *a*.

**Figure 1:**
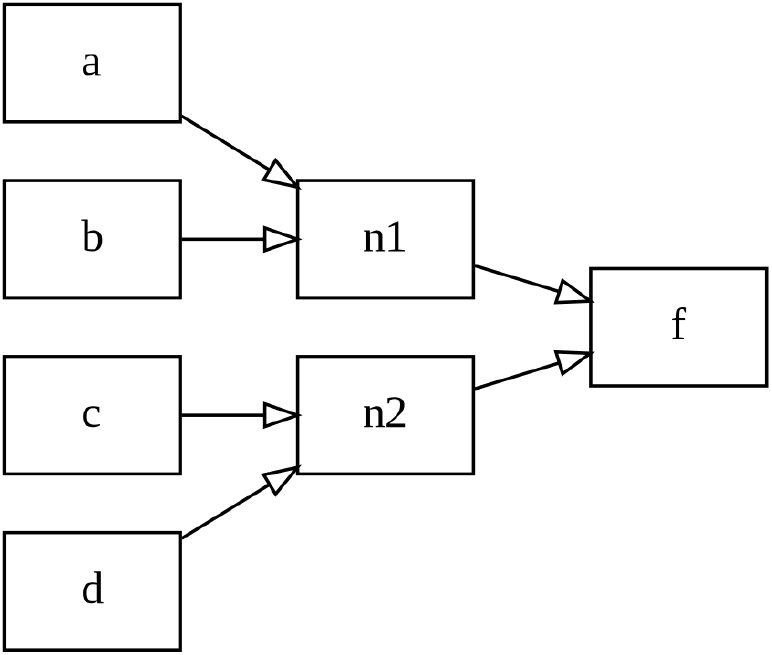
A single elimination tournament with 4 teams.

The number of edges (or arcs) connected to a node is denoted as the degree of the node. The number of arcs pointing *to* a node is called the in-degree, and the number of arcs pointing *away* from a node is called the out-degree.

A phylogenetic tree is a DAG with two types of nodes: tips, which have an in-degree of 0; and inner nodes which have an in-degree of 2. Furthermore, almost all nodes in a phylogenetic tree have an out-degree of 1 and only one dedicated node has an out-degree of 0. This particular node is known as the root.

A set of events with intrinsic time dependencies, (i.e., one event/match must be completed before another event/match) is naturally acyclic. For instance, in a tournament, the winners of the two semi-finals must be determined before the winner of the final can be determined. Since a tournament is also directed, it is a DAG, as outlined in Figures 1 and 2.

**Figure 2:**
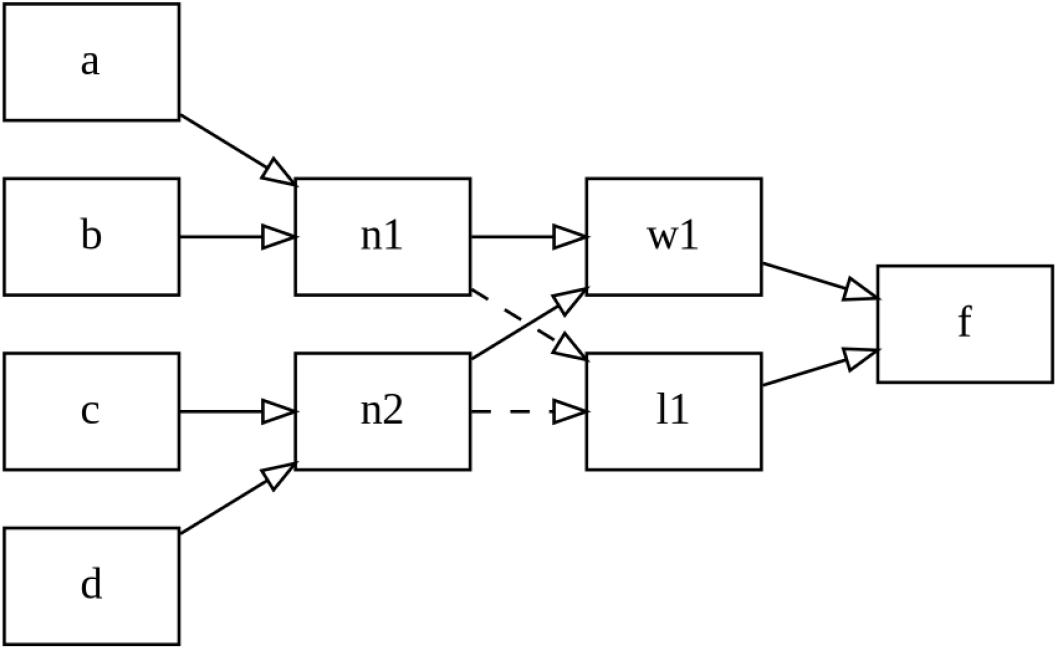
A simple tournament with a losers bracket. The dashed line represents the loser of a specific match. So, in this case, the losers of `n1` and `n2` will play each other in the match `l1`.

In analogy to a phylogenetic tree (for brevity: phylogeny), tournaments have 2 types of nodes: competitors which have an in-degree of 0; and matches, which have an in-degree of 2. However, in contrast to a phylogeny, nodes are allowed to have an out-degree of either 1 or 2. The possible out-degree of 2 is to account for the loser of a match moving down to a losers bracket (also called the lower bracket), albeit tournament matches with an out-degree of 1 appear to be more common. Nonetheless, each tournament retains the special node, or match, with an out-degree of 0. This is the final match that will yield the winner of the tournament.

The above difference between phylogenies and tournaments introduces a complication. When computing the probability of a specific winner for a particular match, we must account for all possible paths that could have lead the specific winning team to this particular match. Consider the example provided in Figure 2. Here, competitor a can either arrive at match `f` via match `w1` or via `l1`. Thus, in order to accurately compute the probability of competitor a winning match `f`, we need to sum over the probabilities of arriving at `f` via `w1` or `l1`.

Fortunately, if we desire to account for these multiple possible paths, we only need to consider the two matches immediately preceding any given match. However, we need to assume that the probability of winning a match is ‘path independent’. This assumption allows us to ‘forget’ about the previous matches that a competitor has played and restricts the calculation to the match at hand. Please see Section 3.1 for a comprehensive description of the mathematical details.

## 3 Method

Initially, we discuss the theory of computing the winner distribution for a single match in Section 3.1. Subsequently, we discuss how we use this theory to efficiently compute the distribution of winners for a general tournament in Section 3.2. We describe the software tool that implements this method in Section 3.3. Finally, we outline how we performed our predictions for the UEFA Euro 2020 European Football Championship in Section 3.4.

### 3.1 Theory

Initially, we provide some definitions. The win probability vector (WPV) for a given node in the tournament tree is a vector containing the probabilities of observing a given team at that node. We denote the probability of team *a* winning over team *b* in a single match as:

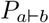

that is, the probability that ‘team *a* beats team *b*’. As such, a *general* WPV has *n* entries, where *n* is the number of teams in the knock-out tournament.

Suppose that we have the most simple tournament with only two teams, *a* and *b*. Then, the WPV which describes this tournament is:

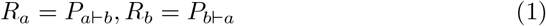

Because this constitutes a trivial case, the calculation is straight-forward. To be able to extend this to non-trivial cases, we will artificially complicate the above expression. First, we introduce the WPVs for *a* and *b* as *w* and *y*. Since *a* and *b* are ‘tips’ of the tree, we can set the probability of observing the team at that node to 1.0 for the team, and 0.0 for all other teams. By doing so, we obtain the expression

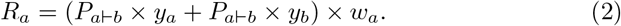

Further, we define *P_t⊢t_* := 0.0 for any team *t*. So, because *P_a⊢a_* := 0.0 and *y_b_* := 1.0, we can reduce Equation 2 to Equation 1. Using this property, we can construct a general expression for the WPV at any particular node of a tournament (including the final) with previous matches already computed as

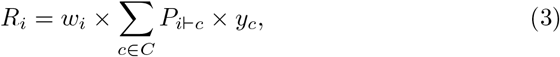

where *R_i_* is the *i*-th entry of the WPV, and *C* is the set of competitors. For multi-elimination tournaments, we also need to account for the fact that a competitor *c* ∈ *C* can come from both sides of the tournament. Therefore, we need to include a second term in the expression to accommodate the other side:

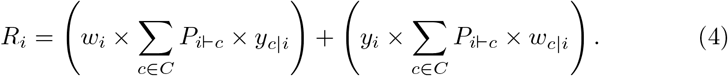

We calculate *w_c|i_ = w_c_/*(1 − *w_i_*). We interpret this as the probability of observing competitor *c* at *w* given that competitor *i* is the opponent in the match. Thus, Equation 4 is the full general expression for the WPV of a multi-elimination tournament. The final complication is that *P_a⊢b_* might be a ‘best of *k*’ series of play-off matches (e.g., in the National Basketball Association (NBA) playoffs). This *k* can also vary over the duration of the tournament since early matches are often ‘best of 1’ with *k* := 1, whereas later matches might be ‘best of 5’ with *k* := 5. We can account for this by introducing a new *P^′^* which represents the pairwise probability of winning the ‘best of *k*’.

### 3.2 Implementation

In order to compute the most likely winner of the entire tournament, we need to compute the WPV for the final match at the root of the tree. For example, in Figure 1, the match `f` must be evaluated. However, in order to compute this, the corresponding WPVs for matches `n1` and `n2` must be computed, as these represent the intermediate results used in Equation 4. Analogously, for the tournament in Figure 2, the WPVs for matches `n1` and `n2` must be evaluated before the WPV for either match `w1` or `l1` can be computed.

Therefore, the tournament matches must be evaluated in the correct temporal order to yield a valid result. This sequence of operations on a tree is analogous to how a likelihood score is computed on a phylogeny. As outlined before, the key difference is that a competitor might be able to traverse multiple paths to reach the final match. Instead of finding a simple traversal, we need to find a topological sorting of the tournament DAG. A topological sorting is a list of the nodes of a DAG such that, if the list is read from left to right, all dependencies are satisfied. Note that, all DAGs can be sorted topologically [8]. If the DAG is a simple binary tournament tree or phylogenetic tree, a topological sorting can easily be obtained via a post order traversal of the tree. In other words, we can calculate the WPV of the final by computing and storing WPVs bottom up at every node, starting from the leaves/tips of the tree and moving toward its root (the final). This procedure is analogous to the computation of the so-called Conditional Likelihood Vectors (CLVs) on phylogenetic trees via the Felsenstein pruning algorithm.

The most important detail missing is how to obtain the pairwise win probabilities *P*. In the preceding Section 3.1, we intentionally considered these probabilities as black boxes for the following two reasons. First, there exist many possible and sophisticated ways to compute *P* as described, for instance, in recent work by Groll *et al.* [5] or in the classic paper by Dixon and Coles [2]. All approaches exhibit advantages as well as disadvantages. Second, computing these probabilities is not the main contribution of this work as we focus on (i) the similarity between phylogenetics and tournaments and (ii) the amount of computations that we can save by applying the Felsenstein pruning algorithm to efficiently and exactly calculate tournament win probabilities, given *P*.

The complexity of a naïve evaluation amounts to 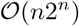 floating point operations. In contrast, the complexity of a Phylourny-like method is 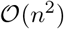. We provide the underlying idea for the time complexity in the following. Consider the probability that competitor 1 wins in an *n* := 8 competitor single elimination tournament. A *single* term for *just one team* is

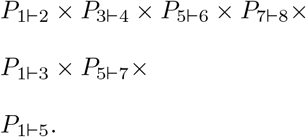

We have organized the above term into layers, one for each ‘tier’ of the tournament. Since we start with *n/*2 matches, and halve their number every time, we have a known series of matches which sum to *n* − 1. Now, to count the number of terms, we note that we have a ‘choice’ for every factor that does not involve competitor 1. For example, we also need to compute the term where competitor 4 beats competitor 3. This means that there are 2*^n^* − log(*n*) terms. If we combine these, we obtain the total expression 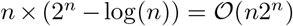.

The naïve space complexity can also be derived from this example. The space required to compute this expression comprises the table of pairwise probabilities, and two additional floating point values. One floating point values is used for the running total of the probability, and the other is used to compute the current term. Therefore, the total space requirement is 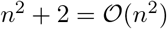.

The time complexity of a Phylourny-like method is more straight-forward to compute. As we can reuse intermediate results from each match, we solely need to evaluate the WPV for each match once. Additionally, there are *n* elements in the WPV. Thus, we need to compute *n* values per 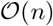 matches. This yields a time and space complexity of 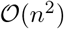. The space complexity is the same for the naïve method and the Phylourny method as the pairwise win probabilities must be stored.

### 3.3 Software

A `C++` reference implementation of our algorithm is available on GitHub^3^ under GNU GPL version 3.0. The software only requires `CMake` to build and also requires `git` to download. We used this implementation (version v0.1.0) to compute the EURO 2020 predictions presented in Section 3.4.

### 3.4 Prediction of the UEFA EURO 2020

In order to predict the winner of the UEFA EURO 2020, we need to estimate the pairwise win probabilities of the competing national teams which constitutes a challenging task. However, as we are only interested in verifying our method for *tournaments* we can, for instance, use the match history from the group stages. Therefore, our prediction for the winner of the championship was conducted *after* the group stage, but *before* the knockout stage.

Nonetheless, using the match history from the group stage and omitting draws implies that the data are sparse. For instance, for the UEFA EURO 2020 tournament, we could only use the results of 10 matches from the group stage. As many teams will not play each other before the elimination stage, the estimation of pairwise win probabilities therefore remains difficult. To overcome this challenge, we deploy two methods. First, we perform a Bayesian sampling of plausible pairwise win probabilities, given the data from the group stage. Second, we utilize the predictive power of existing expert models to infer pairwise win probabilities, which are subsequently used to predict a winner.

#### 3.4.1 MCMC Sampling

The Bayesian sampling is performed via a Markov Chain Monte Carlo (MCMC) search. At each MCMC step, a pairwise win probability matrix is proposed, and the associated WPV is computed for the tournament. Additionally, the likelihood of the pairwise win probability matrix (*P* in earlier sections) is computed using the match data from the group stage. This likelihood represents how likely the proposed pairwise win probability matrix is, given the match history data. This likelihood is different from the phylogenetic likelihood mentioned previously. Informally, a more likely pairwise win probability matrix is one which better explains or fits the previous match history.

The MCMC sampling procedure should be continued until the chain has reached ‘apparent convergence’. Note that true convergence can only be attained if the MCMC sampling is executed infinitely. Further, only the lack of convergence can be assessed via appropriate tools. Hence, as assessing the convergence of MCMC is known to be difficult, we only draw a fixed number of samples. However, computing a single sample using Phylourny is trivial. Therefore, we are able to compute a very large number of samples in a moderate amount of time. For a *n* := 16 competitor single elimination tournament, we were able to evaluate 10 million samples in approximately 5 minutes using a high end 2000 EUR laptop. Therefore, predictions for the UEFA 2020 knock-out stage were performed using 10 million samples. This corresponds to approximately 33, 333 exact calculations of the tournament final WPV per second. We believe that using 10 million samples is justified, as the state space for *this* specific tournament is not excessively large, and should be sufficiently sampled with this number of samples.

Our MCMC search is straight forward. Each *P_a⊢b_* is proposed according to a uniform prior, with *P_b⊢a_* = 1 − *P_a⊢b_* for all competitors *a* and *b*. We sample every proposal, and we run the search until we have obtained 10 millions samples. When computing the summary statistics, we discard the first one million samples as burn in.

Once we have obtained all samples from the MCMC procedure, we can compute two predictions: the maximum likelihood prediction (MLP), or the maximum marginal posterior prediction (MMPP). The MLP is simply the prediction given by the pairwise win probability matrix with the highest likelihood score, whereas the MMPP is the average prediction from all samples. Because an MCMC search will sample the posterior with a probability distribution hopefully approximating the true posterior, the average over all samples is approximately the average of the posterior. The difference between these two predictions is one of philosophical nature rather than mathematics, as they encapsulate distinct interpretations about what ‘really’ matters. The school of thought advocating the MLP claims that the only thing that matters is the *most likely* outcome, regardless of the underlying distribution, whereas the school of thought supporting the MMPP claims that the *totality of evidence* is what matters. A discussion about the merits of these two schools of thought is beyond the scope of this paper.

#### 3.4.2 Model Based Forecast

To perform a model based prediction, we use an existing model (we call this the Lazy Method (LM)), published by Groll *et al.* [5], who have also published previous football tournament predictions, for instance, for the Woman’s World cup in 2019 [6]. Groll *et al.* deploy a random forest approach, utilizing match histories, bookmaker odds, and average player ratings to obtain a pairwise win probability matrix as well as general predictions for the UEFA EURO 2020 tournament. We have thus used their pairwise win probability matrix which was published on the web^4^, and used Phylourny to compute the WPV of the tournament.

The pairwise win probabilities from LM are input directly to Phylourny and used to compute the WPV. We performed no modifications to the data, other than to remove the teams that do not participate at the knock-out stage. This *P* matrix is the one we trust most due to the broad input data from distinct sources being used and the tournament prediction track record of the associated research group.

## 4 Results

Overall, we computed 3 predictions for the three alternative pairwise win probability calculations: Lazy Method (LM), Maximum Likelihood Prediction (MLP), and Maximum Marginal Posterior Prediction (MMPP). Phylourny was executed as follows to calculate the predictions:

> `./phylourny --teams euro2020.ini --matches euro-match-history.csv --probs euro.probs.csv --prefix EURO2020`

The respective input data and relevant output files of Phylourny are available at https://cme.h-its.org/exelixis/resource/download/phylourney-data.tar.bz2.

The predicted tournament winner and loser of the final for each of the three methods is summarized in Table 1.

**Table 1:** Table of pairwise win prediction methods, the team winning the tournament and the team losing the final induced by the respective method, and the actual result.

| Prediction Method | Winning Team | Losing Team |
| --- | --- | --- |
| LM | England | France |
| MLP | Netherlands | France |
| MMPP | Netherlands | Italy |
| Actual Result | TO BE INSERTED on July 11 |  |

**Table 2:** Complete WPVs for the knock-out phase of the UEFA Euro 2020 comprising all 16 teams using the LM, MLP, and MMPP methods for predicting pairwise win probabilities. The two teams with the highest probability of winning the tournament per method are shown in bold font.

| Pred. Meth. | Bel | Por | Ita | Aus | Fra | Swi | Cro | Spa | Swe | Ukr | Eng | Ger | Net | Cze | Wal | Den |
| --- | --- | --- | --- | --- | --- | --- | --- | --- | --- | --- | --- | --- | --- | --- | --- | --- |
| LM | 0.057 | 0.100 | 0.076 | 0.011 | **0.149** | 0.016 | 0.025 | 0.102 | 0.018 | 0.006 | **0.165** | 0.114 | 0.077 | 0.007 | 0.011 | 0.064 |
| MLP | 0.100 | 0.004 | 0.159 | 0.000 | **0.192** | 0.002 | 0.001 | 0.094 | 0.089 | 0.000 | 0.103 | 0.025 | **0.225** | 0.000 | 0.001 | 0.004 |
| MMPP | 0.074 | 0.047 | 0.080 | 0.058 | 0.077 | 0.050 | 0.052 | 0.064 | 0.066 | 0.041 | **0.083** | 0.061 | **0.088** | 0.050 | 0.054 | 0.053 |

Beyond the discrete predicted winner, the scientifically more interesting result is the full WPV for LM, MLP, and MMPP as predicting football matches is know to be notoriously difficult because of the low number of goals being scored that induces a substantial impact of chance onto the final result [7]. To this end, football match predictions do exhibit a high degree of uncertainty and winner predictions should thus be displayed as per-team probabilities, that is, as WPVs. We include the respective WPVs under LM, MLP, and MMPP in Table 4. The two teams with the highest probability of winning the tournament per pairwise prediction method/matrix *P* are shown in bold font. These data can be used to assess the prediction accuracy of our method in retrospect, once the tournament is over on July 11, 2021.

Comparing our prediction using the LM method with that of the original paper describing the LM method [5] which we denote as oLM we obtain: 16.5% for England (oLM: 13.5%), 14.9% for France (oLM: 14.8%), 10.2% for Spain (oLM: 12.3%), 10.0% for Portugal (oLM: 10.1%), and 11.4% for Germany (oLM: 10.1%). The slight deviations in the predictions despite using the exact same *P* matrix are due to the fact that the oLM values were computed *before* the group phase including the prediction of the by then still *unknown* tournament tree for the elimination phase. In contrast to this, our predictions were computed *after* the group phase for a *known* tournament tree.

## 5 Conclusion

We have shown that the problem of predicting tournament winners is sufficiently similar to phylogenetic likelihood calculations such that analogous computational techniques can be applied. We have demonstrated this by developing methods inspired by computational phylogenetics to predict tournaments, and that applying these methods yields substantial computational speedups in terms of theoretical run time complexity. In addition, we can calculate the final WPV of a tournament exactly instead of using simulations to approximate it. This also allows, for instance, for a seamless deployment of MCMC methods such as illustrated by our admittedly very simple example in Section 3.4.1.

Furthermore, we demonstrate the practicality of these new methods by implementing them into a new software tool called Phylourny.

As we are writing this *before* the tournament enters its knock-out phase, we do not know how successful our method will be at predicting the true outcome^5^. Nonetheless, we can already discuss the two theoretical shortcomings of our approach regardless of the success of our prediction. First, the prediction ‘difficulty’ is predominantly deferred into estimating the pairwise win probability matrix. This constitutes the central problem of tournament prediction, which we do intentionally not directly address. Betting companies with their substantial resources and other researchers have already addressed this problem to a large extent [9]. Instead, we present a *computational* method, which will accelerate the exact computation of final win probabilities, given some estimation of pairwise win probabilities, and a surprising connection between two seemingly unrelated branches of science.

Second, the assumption of path independence might not be true, as competitors might suffer from fatigue from competing in more matches, if a team must proceed through the lower bracket in order to reach the finals or by having to play harder opponents or play over-time. Furthermore, other ‘intangibles’, such as moral or confidence, are hard to quantify, also questioning the path independence assumption. Nonetheless, this path dependence can be addressed via a more involved method of calculating the pairwise win rate matrix, as one can also deploy a match-dependant *P* matrix.

Despite the two deficiencies mentioned above, we have shown that we can compute, both exactly and efficiently, the WPV for a tournament. This is important because, many advanced methods of analysis require exact results to be applicable. For example, when sampling from a posterior using an MCMC search, it is desirable to have an accurate result for each sample. While a sufficient degree of accuracy can be obtained via an appropriately large number of simulations, this approach is computationally expensive and might even become prohibitive. We demonstrate that we can efficiently conduct such an analysis by implementing our own (naïve) MCMC analysis of the UEFA EURO 2020 football tournament.

While we consider this work as being complete, there exist further areas of investigation that can be explored. An example is exploring the ‘stability’ of complicated tournaments by slightly perturbing *P* and examining the resulting probabilistic outcome. Due to the increased computational efficiency and the ability of Phylourny to *exactly* calculate the final WPV, such studies are substantially more tractable now. Another area of interest would be to further develop the MCMC sampling. We currently use an extremely naïve MCMC search that could become more efficient by specifying more elaborate methods for proposing new parameters.

## Footnotes

1 A multi-elimination tournament is any tournament where a competitor must loose more than once to be eliminated from the tournament. These are almost always double-elimination tournaments, but one can imagine triple or more elimination tournaments. We use this term in order to highlight the more general nature of this method

2 The 2020 European Football Championship was postponed to summer 2021 due to the COVID-19 pandemic. This how we can predict a tournament in 2020 with a paper written in 2021.

3 https://github.com/computations/phylourny

4 https://www.zeileis.org/news/euro2020/

